# Expression of *Brassica napus* GLO1 is sufficient to breakdown artificial self-incompatibility in *Arabidopsis thaliana*

**DOI:** 10.1101/2020.04.27.064394

**Authors:** Patrick Kenney, Subramanian Sankaranarayanan, Michael Balogh, Emily Indriolo

**Author notes:** Corresponding author: Emily Indriolo.

## Abstract

Members of the Brassicaceae family have the ability to regulate pollination events occurring on the stigma surface. In *Brassica* species, self-pollination leads to an allele specific interaction between the pollen small cysteine-rich peptide ligand (SCR/SP11) and the stigmatic S-receptor kinase (SRK) that activates the E3 ubiquitin ligase ARC1 (Armadillo repeat-containing 1), resulting in proteasomal degradation of various compatibility factors including Glyoxalase I (GLO1) which is necessary for successful pollination. Suppression of GLO1 was sufficient to reduce compatibility, and overexpression of GLO1 in self-incompatible *Brassica napus* stigmas resulted in partial breakdown of the self-incompatibility response. Here, we verified if *Bn*GLO1 could function as a compatibility factor in the artificial self-incompatibility system of *Arabidopsis thaliana* expressing *Al*SCR_b_, *Al*SRK_b_ and *Al*ARC1 proteins from *A. lyrata*. Overexpression of *Bn*GLO1 is sufficient to breakdown self-incompatibility response in *A. thaliana* stigmas, suggesting that GLO1 functions as an inter-species compatibility factor. Therefore, GLO1 has an indisputable role as a compatibility factor in the stigma in regulating pollen attachment and pollen tube growth. Lastly, this study demonstrates the usefulness of an artificial self-incompatibility system in *A. thaliana* for interspecific self-incompatibility studies.

## Introduction

Many lineages of flowering plants have developed self-incompatibility systems to prevent inbreeding and encourage genetic diversity through outcrossing (Charlesworth and Vekemans 2005; de Nettancourt 2001; Hiscock and Allen 2008; Iwano and Takayama 2012). In the mustard family of plants, which includes the model dicot, *Arabidopsis thaliana*, *Brassica napus* (canola) and other economically important species of *Brassica*, most of the species have a self-incompatibility system to reject ‘self’ pollen. In the Brassicaceae, the self-incompatibility system is sporophytic, where the recognition occurs between the diploid male and female parts of the plant. Acceptance is mediated by the stigma, a dry type, which will only allow the transfer of water from the stigmatic papillae cells to the desiccated pollen grain if it is compatible (Heslop-Harrison 1979). A rejected ‘self’ pollen grain will be unable to hydrate and germinate on the stigma surface as a result of the activation of the self-incompatibility response. One of the major exceptions is *A. thaliana*, a species that has lost its self-incompatibility system (Boggs et al. 2009; Castric et al. 2014; Indriolo et al. 2012; Schierup et al. 2006; Tsuchimatsu et al. 2010). Thus, the study of this self-incompatibility system has been predominantly focused on *Brassica spp* and *Arabidopsis lyrata* (Indriolo et al. 2012; Ivanov et al. 2010; Iwano and Takayama 2012; Kusaba et al. 2001; Schierup et al. 2006; Tantikanjana et al. 2010).

Through the analysis of *Brassica spp*, the critical components of this self-incompatibility (SI) signaling pathway have been determined. These include the pollen coat ligand Small Cysteine Rich, SCR, or S-locus Protein 11, SP11 (referred to as SCR from here on), the female receptor kinase S-locus Receptor Like Kinase (SRK) which can recognize SCR (Kachroo et al. 2001; Shimosato et al. 2007; Tantikanjana et al. 2010). SRK becomes phosphorylated and activated, and with the cytoplasmic receptor like kinase, M-locus Protein Kinase (MLPK), these kinases phosphorylate and activate downstream signaling components (Kakita et al. 2007; Murase et al. 2004; Samuel et al. 2008). The Armadillo Repeat Containing 1 (ARC1) E3 ubiquitin ligase is one of the proteins phosphorylated and activated by SRK and MLPK in this signaling pathway (Gu et al. 1998; Stone et al. 2003; Stone et al. 1999). ARC1 has been shown to target several compatible pollination factors that are involved in the basal compatibility response including, EXO70A1, Phospholipase D1, (PLD1) and *B. napus* Glyoxalase 1 (*Bn*GLO1) for ubiquitination (Samuel et al. 2009; Sankaranarayanan et al. 2017; Sankaranarayanan et al. 2015; Scandola and Samuel 2019). EXO70A1, a member of the exocyst complex, mediates the delivery of multi-vesicular bodies (in *Brassica spp*) and secretory vesicles (in *A. lyrata* and *A. thaliana*) to the pollen contact point that result in pollen grain hydration (Indriolo et al. 2014; Safavian and Goring 2013; Safavian et al. 2015; Samuel et al. 2009). PLD1 activity during compatible pollination promotes vesicle fusion at the membrane, facilitating exocytosis that is essential for pollen germination (Scandola and Samuel 2019). Glyoxalases (GLO) are enzymes involved in detoxification of methylglyoxal (MG), a cytotoxic byproduct of glycolysis (Sankaranarayanan et al. 2017). *Bn*GLO1 prevents the buildup of cytotoxic methylglyoxal (MG) within the stigmatic papilla cells in response to a ‘self’ pollen (Sankaranarayanan et al. 2015). *Bn*GLO1 was clearly shown to be a target of ARC1 for ubiquitination, and was more likely to be ubiquitinated when the protein had a MG modification as a part of the increase in MG within the papilla following perception of a ‘self’ pollen grain and the activation of the self-incompatibility signaling pathway (Sankaranarayanan et al. 2015). Therefore, during a compatible pollination, *Bn*GLO1 detoxifies MG levels in the stigma and helps to prevent the MG modification of other proteins. During a self-incompatible pollination, the basal pollen response is blocked, including the MG detoxification by *Bn*GLO1. Overexpression of *Bn*GLO1 in self-incompatible *B. napus* stigmas resulted in the breakdown of the self-incompatibility response, causing the acceptance of ‘self’ pollen (Sankaranarayanan et al. 2015).

Self-incompatibility in *A. lyrata* has been characterized to determine the changes that have occurred between lineage I, including *Arabidopsis spp* and lineage II, which includes *Brassica spp. A. lyrata* was also studied to determine the events that lead to the loss of self-incompatibility in the closely related species, *A. thaliana* (Castric et al. 2014; Indriolo et al. 2012; Nasrallah et al. 2002; Tsuchimatsu et al. 2010). A notable difference from *Brassica spp*. is that MLPK does not appear to play a role in self pollen rejection in *A. lyrata* (Castric et al. 2014; Indriolo et al. 2012; Nasrallah et al. 2002; Tsuchimatsu et al. 2010). RNAi knock down of *ARC1* in *A. lyrata sp. patrea* resulted in plants that were able to accept incompatible pollen grains and demonstrated that *ARC1* is necessary for the rejection of self-pollen in *A. lyrata* (Indriolo et al. 2012). A survey of over 300 ecotypes of *A. thaliana* showed that all of the accessions had the same *ARC1* deletion in their genome indicating that the breakdown of ARC1 in *A. thaliana* predated the breakdown of SCR and SRK (Castric et al. 2014; Indriolo et al. 2012).

SCR, SRK and ARC1 were demonstrated to have conserved roles in *A. lyrata*, as observed in *Brassica spp*. This lead us to the idea that ARC1 could potentially restore a more robust self-incompatibility response in *A. thaliana* through stigma specific expression of the transgene in various ecotypes of *A. thaliana*. Previous studies had reconstructed a self-incompatibility response in *A. thaliana* expressing *AlSCR_b_* and *AlSRK_b_* alone, but there was a wide range of variability in the robustness of the response based on individual ecotype (Boggs et al. 2009; Nasrallah et al. 2004; Strickler et al. 2013; Tsuchimatsu et al. 2010). Through rigorous surveys of *A. thaliana* ecotypes, some have been identified with functional copies of SRK, such as the Wei-1 and Old-1 accessions. The Wei-1 plants have a functional *SRK_a_* and the *SCR_a_* gene is inverted; when the inversion was returned to its original state, transgenic Wei-1 plants were able to reject ‘self’ pollen with the a-haplotype (Tsuchimatsu et al. 2010). With this variation in *A. thaliana* ecotypes in regards to the retention of functional SRKs and the different phenotypes observed in transgenic *A. thaliana* plants expressing *AlSCR_b_* and *AlSRK_b_*, experiments were performed to look at the role of ARC1 in an artificial transgenic self-incompatibility system. Therefore, two *A. thaliana* ecotypes were selected: Col-0, which in general did not have a strong self-incompatible phenotype with *AlSCR_b_* and *AlSRK_b_*, and the Sha ecotype which did show a self-incompatible phenotype when expressing *AlSCR_b_* and *AlSRK_b_* (Boggs et al. 2009). In addition to the stigma specific expression of *AlSCR_b_* and *AlSRK_b_*, either *AlARC1* or *BnARC1* were expressed in the stigma under the control of the SLR1 promoter (Indriolo et al. 2014). With the addition of ARC1, a robust self-pollen rejection phenotype was observed in both the Col-0 and Sha ecotypes including aniline blue stains, seed set, and at the ultrastructural level, in the removal of the secretory vesicles likely by autophagy (Indriolo et al. 2014). The generation of self-incompatible *A. thaliana* with a solid and robust phenotype were a result of the addition of *Al*ARC1 or *Bn*ARC1 with *Al*SCR and *Al*SRK; these results indicated ARC1 plays a conserved role between lineage I and lineage II of the Brassicaceae (Indriolo et al. 2014). Furthermore, a yeast two hybrid assay demonstrated that *Bn*ARC1 and *Al*ARC1 could interact with *Al*SRK1 and *Bn*SRK910 showing that these proteins kept the ability to recognize each other as a part of this signaling pathway despite being phylogenetically separated (Indriolo and Goring 2016).

The role of *Bn*GLO1 has been well defined in *B. napus* self-incompatibility. This poses the question if this compatible stigma factor has a role in *Arabidopsis spp*. in the artificial self-incompatibility system of *A. thaliana*. To determine if *Bn*GLO1 plays a role in the acceptance of pollen grains, it was transformed into *A. thaliana* self-incompatibility transgenic lines in the Sha ecotype that contained *Al*SCR_b_, *Al*SRK_b_ and *Al*ARC1. By adding stigma specific expression of *Bn*GLO1 in addition to the endogenous expression, these lines would be similar to a mild over expression line. If the overexpression of *Bn*GLO1 in addition to the endogenous *At*GLX1 (and likely target of *Al*ARC1) was able to break down the self-incompatibility response it would demonstrate that glyoxalase activity is key to accepting compatible pollen.

## Materials and Methods

### Plant Materials

Sha wild-type seeds (CS22652), Sha self-incompatible lines *Al*ARC1+*Al*SRK_b_+*Al*SCR_b_ (Indriolo et al. 2014), and Sha self-incompatible lines (*Al*ARC1+*Al*SRK_b_+*Al*SCR_b_) transformed with *Bn*GLO1 (Sankaranarayanan et al. 2015) were used in this study.

The wild-type Sha seed stocks came from the Arabidopsis Research Center (ABRC), Columbus, Ohio, USA. All transgenic plants were generated as described in Indriolo et. al. 2014 or in this work.

### Plant transformation

All transgenic plants were generated by floral dip as described in Clough and Bent 1997. Sha self-incompatible lines *Al*ARC1+*Al*SRK_b_+*Al*SCR_b_ plants (Indriolo et al. 2014) were dipped in *Agrobacterium tumefaciens* containing the *Bn*GLO1 construct (Sankaranarayanan et al. 2015).

### Genotyping of transgenic lines

The *A. thaliana* Sha *Bn*GLO1+*Al*ARC1+*Al*SRK_b_+*Al*SCR_b_ were genotyped by PCR (primers in Supplemental Table 1, Supplemental Figure 3), to determine transformants that had all three of the plasmids present in the genome. The *Al*ARC1+*Al*SRK_b_+*Al*SCR_b_ plants were siblings of seeds studied in Indriolo et. al. 2014 which had been previously characterized.

### Plant growth conditions

Plants were grown under long day conditions; 16 hours light/8 hours dark with a daytime temperature of 22°C and a nighttime temperature of 17°C. *A. thaliana* seeds were surface sterilized with 1 volume bleach and 2 volumes Triton X-100 for 15 minutes and washed with sterile dH_2_O, 4 times. After sterilization, all seeds were stratified for 6 days at 4°C and were plated out on ½ Murashige & Skoog, pH 5.8 with 0.4% w/v phytoblend agar plates. Once the seedlings had 2 true leaves, they were transferred to soil (ProMix-BX + mycorrhizal species, Quebec, Canada). The relative humidity of the chambers was recorded to be between 35-55% during all experiments.

### Pollen adhesion, pollen tube growth and seed set analysis

Pollen grain adhesion and pollen tube growth were observed on freshly opened flowers. To determine the correct age of the freshly opened flowers, stage 12 flower buds (the final bud stage prior to bud opening; (Smyth 1990) from wild-type Sha and transgenic plants were emasculated and covered in plastic wrap to prevent drying of the tissues overnight. 24 hours later, individual anthers from either Sha wild-type flowers or transgenic plants were used to manually apply pollen grains to the stigmas of the emasculated flowers when the papillae were fully elongated. 2 hours-post pollination, whole pistils were removed from the plant and placed in a fixative solution (300μL of ethanol:glacial acetic acid [3:1]) at room temperature for 30 minutes. Fixative was then removed, and the pistils were washed with sterile dH_2_O three times and then incubated in 300μL of 1N NaOH for 1hr at 60 °C. Next, the 1N NaOH was removed and the pistils were washed with sterile dH_2_O three times and then stained with 300μL of 0.1% (w/v) aniline blue at 4°C overnight.

To visualize pollen grain adherence and pollen tube penetration, whole pistils were mounted on slides with mounting media (4% (w/v) propyl gallate) to prevent photobleaching. Images were captured with a Zeiss Axio-observer fluorescence microscope. At minimum, 10 pistils were imaged per each individual line for pollen grain adherence (Brightfield) and pollen tube growth (DAPI-358/463nm). Self-pollen was used in all experiments to determine acceptance or rejection of the pollen on the stigma. All pollination assays were performed at a relative humidity level of between 35-55% and were monitored with a digital hygrometer in the chamber at all times.

To determine seed set, the number of seeds per silique were tallied for Sha and each independent transgenic line (n=10). To determine pollen grain adherence to stigmas, the total number of pollen grains were tallied for three different images. The seed set data was graphed and analyzed using R-Studio Version 1.1.463 (2009-2018 RStudio, Inc.). The graph was generated using the ggplot2 package. Statistical analysis included a one-way ANOVA with a Tukey’s HSD Post Hoc test from the “agricolae” package. The R script is included in Supplemental Figure 4.

Pollen grain adherence treatments were examined using Welch’s ANOVA after running Levene’s test for homogeneity. For comparisons between genotypes, we used Dunnett’s multiple comparisons test and a Tukey post-hoc test using the SAS for University software package with an *α* =0.05 for a treatment (in this case genotype) to be determined as statistically significant (Cary, NC). To determine pollen grain adherence to stigmas, the total number of pollen grains were tallied for three different images.

### Confocal imaging of stigmas

To verify expression of *Bn*GLO1 in *A. thaliana* stigmas, whole pistils were removed from stage 13 flowers. To observe stigmatic papilla cells of the transgenic *A. thaliana* Sha ecotype expressing *Bn*GLO1-RFP, pistils were removed from the flowers and placed in a 24 well plate with 1mL sterile dH_2_O containing CellMask Deep Red (Thermo Fischer Scientific) diluted 1:1000. Following incubation for 5 minutes, the solution was removed, the pistils were washed three times with sterile dH_2_O before they were mounted on slides for live cell imaging. Samples were imaged on a Dragonfly 550 spinning disk confocal imaging system mounted on an Olympus IX83 Inverted Microscope using a 30x Apochromat silicon objective (NA 1.05) driven by Andor Fusion software. Images were acquired using an iXon 888 EMCCD camera. Bitplaine Imaris (v9.0.2) software were used to compile and process the images. The final image was generated in Adobe Illustrator.

### Expression Profiling & Protein Sequence Analysis

We generated an expression heat map for the *A. thaliana* glyoxalase I gene family using the genevestigator tool (www.genevestigator.com). This analysis examined the relative expression level of the *A. thaliana* glyoxalase I family in shoot, seedling, stigma, mature pollen, flower, pistil and roots (Hruz et al. 2008). We used The Heat Tree Viewer to analyze the glyoxalase gene family expression in several reproductive tissues and pollen tube growth stages through the pistil. Full length protein sequences for the most closely related copies of Glyoxalase 1 to *Bn*GLO1 were identified by BLAST with the *Bn*GLO1 protein sequence as a query against the translated sequences of the *A. thaliana* and *A. lyrata* genomes. A neighbor-joining tree was generated with MEGA X 10.1 software following multiple alignment using Clustal X multiple sequence alignment tool. Multiple alignment of protein sequences of the nearest homologs to *Bn*GLO1 from *A. thaliana* and with *A. lyrata* were performed using Clustal X and viewed in Jalview. Following sequence alignment, the sequences were trimmed from both the N-terminus and C-terminus to focus on the portion of 163 highly conserved amino acids. Highly conserved amino acids are highlighted in shades of dark gray vs light gray for less conserved residues.

## Results

### Phylogenetic analysis reveals the relationships of *Arabidopsis* glyoxalase homologs

The ten most closely related full-length protein sequences from *A. thaliana* and *A. lyrata* were aligned and then used to generate a neighbor-joining tree (Figure 1A). The results show a clear clade where *Bn*GLO1 (magenta box) is sister to Al1g23010 and At1g11840. The next most closely related clade of glyoxalases is that of At1g67280 and Al2g25800, the two other proteins that were selected to be included in the protein consensus alignment analysis (Figure 1B). Overall, the statistical support based on the bootstrap values for most of the tree is robust with *A. thaliana* and *A. lyrata* glyoxalases forming clades from with a single copy of an *A. thaliana* and *A. lyrata* glyoxalase at the end of each branch. Only a single copy of each species was instead placed sister to a pair of glyoxalases and with high bootstrap support of 97 for Al4g27470 as a single copy with no homologue and support of 98 for At1g80160.

**Figure 1.**
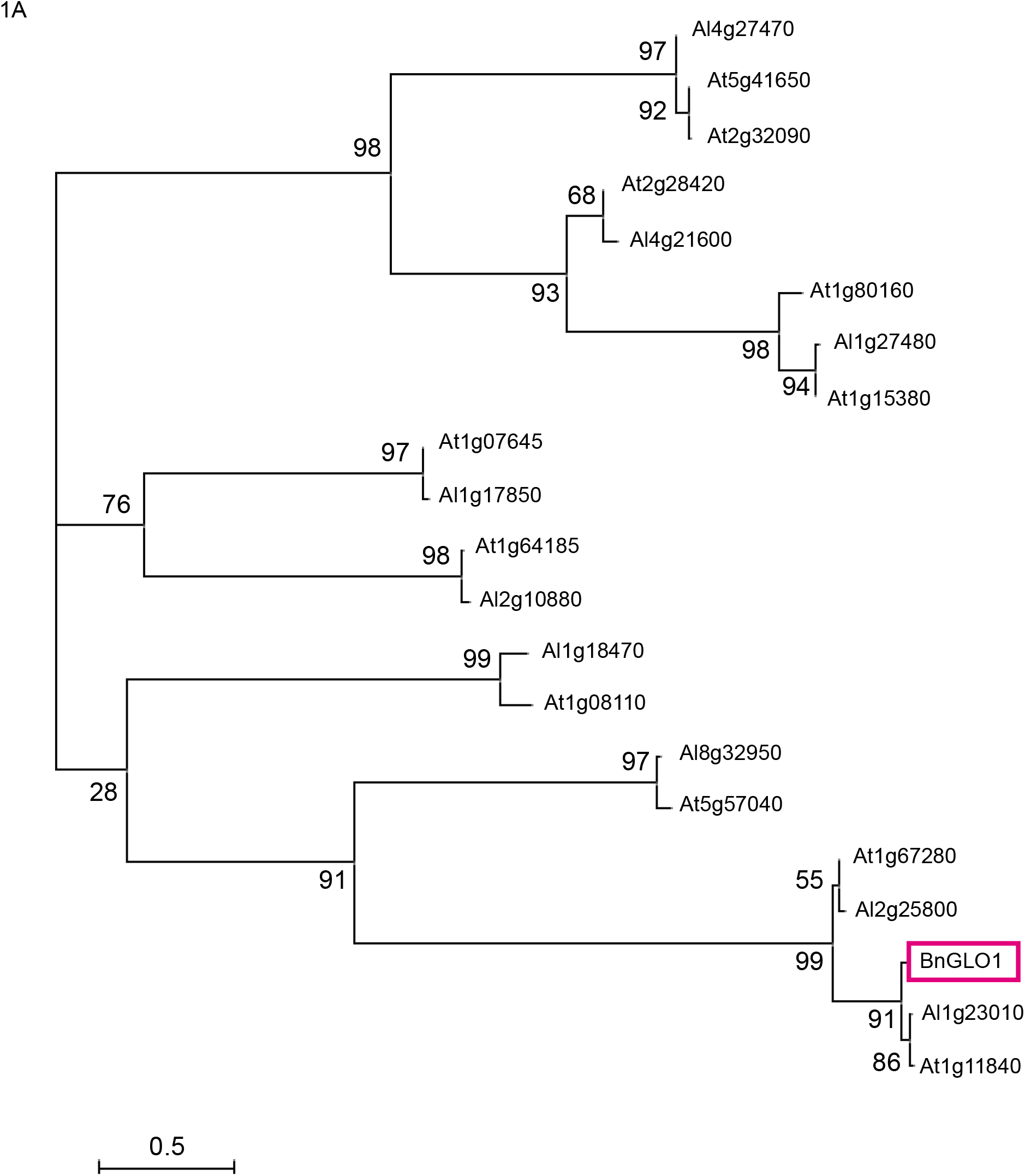
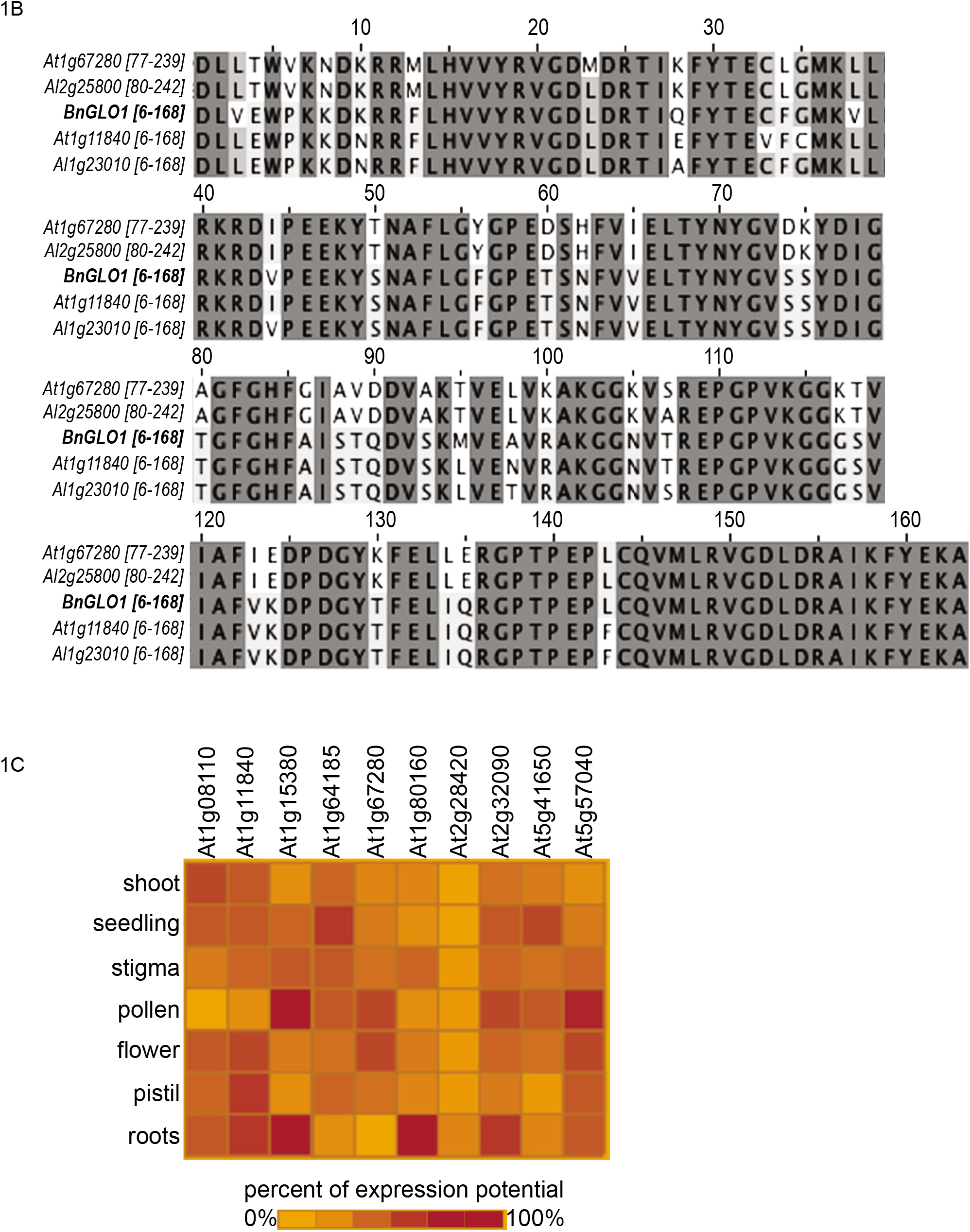
Expression and Alignment of Glyoxalase 1. 1A.) Phylogenetic analysis of the *Bn*GLO1, *Arabidopsis lyrata* glyoxalase homologs and *Arabidopsis thaliana* glyoxalase I family genes. The scale bar represents 0.5 substitutions per amino acid position. Boot strap values are shown at the nodes of each branch of the tree. 1B.) Multiple sequence amino-acid alignment of *Bn*GLO1 with its nearest *Arabidopsis thaliana* and *Arabidopsis lyrata* homologs. Highly conserved amino acids are highlighted in shades of dark gray vs light gray for less conserved residues. Alignment is based on *Bn*GLO1 from residue 6-168. 1C.) Microarray expression data of *Arabidopsis thaliana Glyoxalase 1* family members (www.genevestigator.com). The darkest shade of orange indicates genes at 100% of expression potential while the lightest orange shade indicates 0% of expression potential.

### Protein sequence of BnGLO1 is highly conserved among *Arabidopsis* species

Our driving question was to examine if the overexpression of *Bn*GLO1 would be able to break down self-incompatibility in *A. thaliana* expressing *Al*ARC1-*Al*SRK_b_-*Al*SCR_b_; therefore, we aligned the Glyoxalase 1 protein sequences of *A. thaliana* and *A. lyrata* with that of *Bn*GLO1. This protein alignment (Figure 1B) focuses on the most conserved portion of the full-length protein sequences that correspond to *Bn*GLO1 amino acids 6-168 of the total protein length of 283 amino acids. The protein alignment shows that this region is conserved as the dark grey regions represent 100% conservation between the five protein sequences. More interesting are the regions that show more variation; there are two different conserved domains of the more variable portions of the protein sequences. Of the 24 regions, indicated by the white background, 21 of these regions had an amino acid(s) that are conserved between *Bn*GLO1, At1g11840 and Al1g23010, while a different amino acid was identical in the At1g67280 and Al2g25800 sequences. Examples of these differences are M13 or F13, T50 or S50 and AVD88-90 or STQ88-90. These differences always change relative to *Bn*GLO1, with one *A. thaliana* and *A. lyrata* protein sequence matching *Bn*GLO1 while another *A. thaliana* and *A. lyrata* sequence is not conserved with *Bn*GLO1. These points of variation in the conserved protein sequence indicate that At1g11840 and Al1g23010 are likely working as the functional homologues to *Bn*GLO1 and behave as such in each species in respect to protein activity. By using both protein sequence alignment and phylogenetic analysis, we can conclude that *Bn*GLO1 is likely functional similar to that of At1g11840 and Al1g23010.

### Expression of Glyoxalase 1 in *A. thaliana* tissues

The expression profile of the most closely related glyoxalases to *Bn*GLO1 were examined in a variety of *A. thaliana* tissues. Figure 1C shows the results of the expression analysis; each glyoxalase is expressed as the percent of expression potential across tissues; shoot, seedling, stigma, pollen, flower, pistil and root. The darker the shade of orange, the higher percent of potential expression (maximum 100%). Supplemental Figure 2 shows the expression analysis of glyoxalase 1 gene family members in various reproductive tissues including their expression in pistil after various time points of pollination. It is clear that some glyoxalases have a high level of expression in a wide range of tissues such as *At1g11840*, while *At2g28420* has a very low level of expression potential in all tissues examined (Figure 1C). *At1g11840* has fairly robust expression in the stigma, pistil and flower, and reduced expression in mature pollen (Figure 1C). More importantly when compared to the others, this glyoxalase has the closest relation to *BnGLO1* and is highly expressed in the stigma and pistil before and after pollination. This indicates a putative role in pollen-pistil interactions (Supplemental Figure 2). Other members that might have a role in pollen-pistil interactions include At1g67280 and At1g08110. These genes show a moderate level of expression in the stigma and pistils following pollination (Supplemental Figure 2).

Few glyoxalases like *At1g15380, At1g80160, At2g3209* and *At5g7040* were expressed in the pollen and pollen tubes, indicating that they likely play a role in pollen maturation, pollen tube growth and ultimately in pollen-pistil interactions (Supplemental Figure 2).

### Expression of *BnGLO1* is stigma specific in transgenic lines

Three independently transformed lines of Sha *Bn*GLO1+*Al*ARC1+*Al*SRK_b_+*Al*SCR_b_ were generated by floral dip. All of these lines were first genotyped for the presence of each construct before further characterization (Supplemental Figure 1 & 3; Primers listed in Table 1). Only individuals with all of the transgenes were used for further analysis of self-incompatible pollinations compared to Sha wild-type compatible pollinations.

To verify that the *Bn*GLO1-RFP was expressed in the stigma, unpollinated flowers were imaged using confocal microscopy. To visualize individual papillae, Deep Red CellMask was used to stain the plasma membrane (Figure 2; purple fluorescence). This was used to orient the papillar cells for localization of *Bn*GLO1-RFP in the cells. As previously characterized in *B. napus* and *A. thaliana*, the *Bn*GLO1-RFP (Figure 2; red fluorescence) under the SLR1 promoter was expressed in the stigma and it was found both on the plasma membrane and in the cytoplasm (Sankaranarayanan et al. 2017; Sankaranarayanan et al. 2015). These results demonstrate that the transformation and the expression of *Bn*GLO1-RFP into the self-incompatible *A. thaliana* lines was successful.

**Figure 2.**
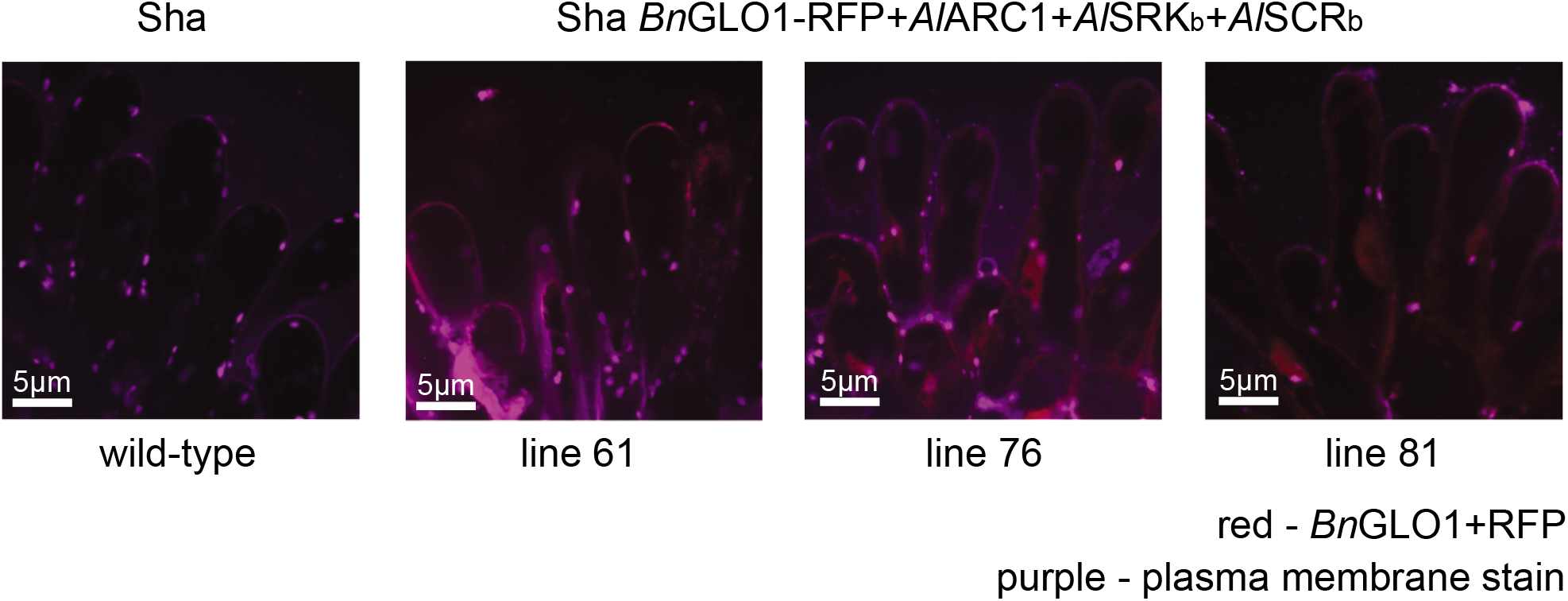
Localization of BnGLO1-RFP in stigmatic papillae. Live cell imaging of stage 13 flowers of *A. thaliana*. Sha wild-type stigmas were stained with the CellMask Deep Red stain to visualize the plasma membrane as shown in purple. Sha *Bn*GLO1-RFP+*Al*ARC1+*Al*SRK*_b_*+ *Al*SCR*_b_* transgenic lines imaged by confocal to visualize *Bn*GLO1-RFP and CellMask Deep Red stain. *Bn*GLO1-RFP is shown in red with the CellMask Deep Red stain in purple for the plasma membrane. Scale bars = 5μm.

### Self-incompatible lines accept compatible pollen with the overexpression of *BnGLO1*

To determine the early pollen response in the Sha *Bn*GLO1+*Al*ARC1+*Al*SRK_b_+*Al*SCR_b_ plants compared to Sha wild-type and self-incompatible Sha *Al*ARC1+ *Al*SRK_b_+*Al*SCR_b_, aniline blue stains were performed to qualitatively examine pollen grain acceptance in the transgenic stigmas. Wild type self-pollinated stigmas (Figure 3A, Supplemental Figure 1) show pollen grain acceptance and ample pollen tube germination and growth 2 hours post pollination. As previously demonstrated, the Sha *Al*ARC1+ *Al*SRK_b_+*Al*SCR_b_ self-incompatible lines (Fig 3A 1-SI, 2-SI and 5-SI, Supplemental Figure 1) showed a strong self-pollen grain rejection phenotype (Indriolo et al. 2014). Few pollen grains were able to adhere to the stigma and very little pollen tube germination and growth were observed for all three self-incompatible lines. In direct contrast, Sha *Bn*GLO1+*Al*ARC1+ *Al*SRK_b_+*Al*SCR_b_ transgenic lines (Fig 3A, 61-SC, 76-SC, 81-SC) appear to be similar to compatible pollen grain acceptance as observed in Sha wild-type stigmas. Pollen tubes germinate readily and penetrate easily at 2 hours after pollination (Fig 3A, 61-SC, 76-SC, 81-SC Supplemental Fig. 2). Therefore, these images indicate that the addition of *Bn*GLO1 in addition to endogenous *A. thaliana* glyoxalases breaks down the self-incompatibly response.

**Figure 3.**
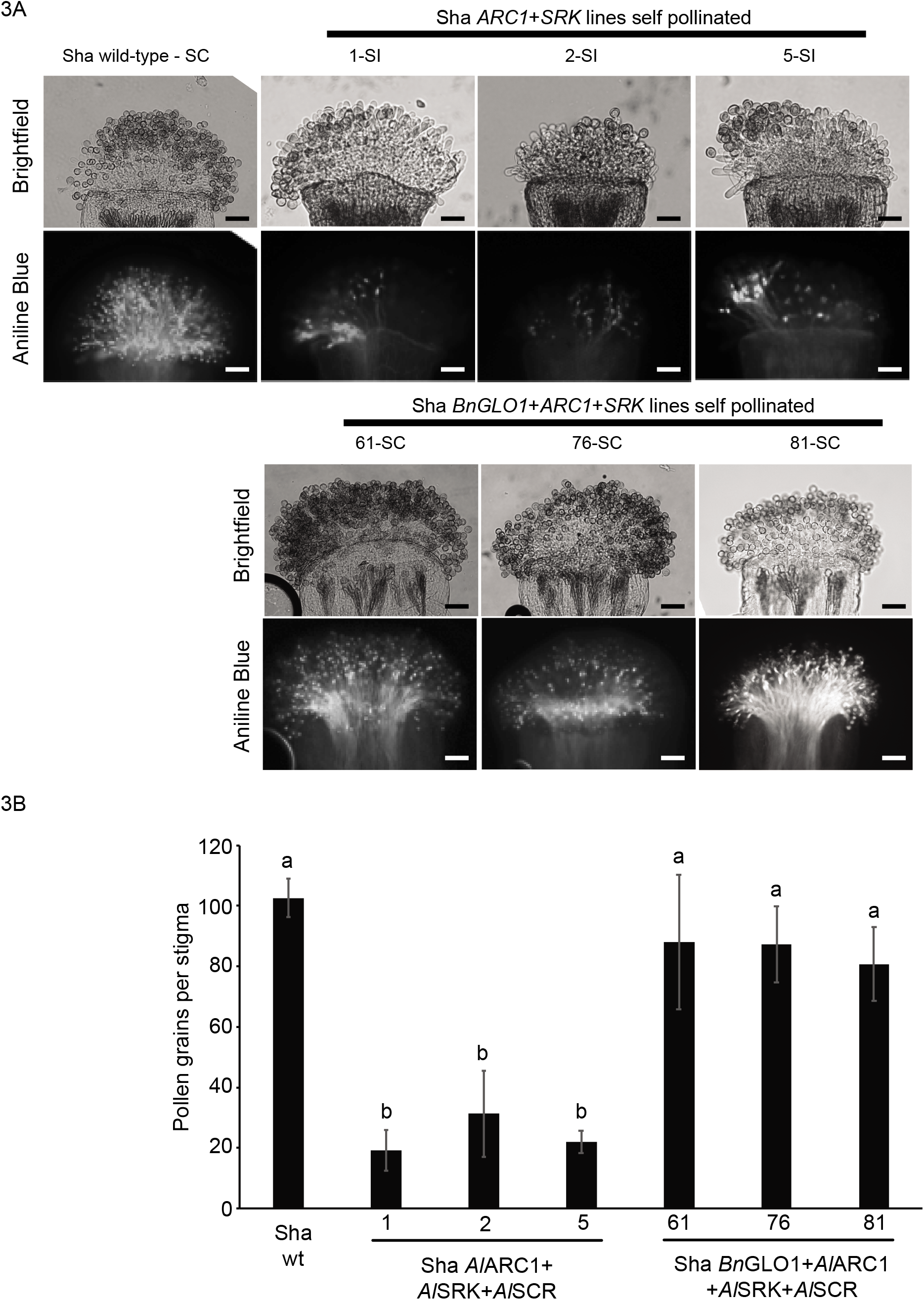
Self-pollination phenotypes of Sha AlARC1+AlSRK_b_+AlSCR_b_ and Sha BnGLO1-RFP+AlARC1+AlSRK_b_+AlSCR_b_ stigmas. 3A.) Brightfield and Aniline Blue stains of stigmas. Sha wild-type self-pollinations show the acceptance of self-pollen grains 2 hours post pollination an example of a self-compatible pollination. Scale bar = 50μm. 3B.) Average number of pollen grains per stigma 2 hours postpollination. Pollen grain adherence was compared by a one-way ANOVA followed by a Tukey post hoc test (p < 0.5), n = 3 stigmas per each transgenic line. Relative Humidity was between 35-55% during the pollinations.

### Self-incompatible lines with *Bn*GLO1 have a compatible pollen response

To quantitatively characterize the breakdown of the self-incompatibility response, pollen grain adherence and seed set were measured in self-pollinations. Pollen grain adherence clearly demonstrated that the average number of self-pollen grains that were able to adhere to the stigma were statistically different between wild-type compatible pollinations, self-incompatible pollinations and self-incompatible pollinations with *Bn*GLO1 (Figure 3B). Wild-type Sha compatible pollen grain adherence was an average of 102.6 ± 6.4 pollen grains per stigma and the self-incompatible lines low pollen grain adherence as previously observed (Figure 3B, Indriolo 2014). Sha *Al*ARC1+*Al*SRK_b_+*Al*SCR_b_ line 1 had on average 19.2 ± 6.7, Sha *Al*ARC1+ *Al*SRK_b_+*Al*SCR_b_ line 2 had on average 31.3 ± 14.2 and Sha *Al*ARC1+*Al*SRK_b_+*Al*SCR_b_ line 5 had on average 22.1 ± 3.7 pollen grains adhered per stigma. Statistically similar to Sha wild-type, Sha *Bn*GLO1+*Al*ARC1+*Al*SRK_b_+*Al*SCR_b_ line 61 had on average 88.0 pollen grains ± 22.3, Sha *Bn*GLO1+*Al*ARC1+*Al*SRK_b_+*Al*SCR_b_ line 76 had on average 87.3 ± 12.6 and the Sha *Bn*GLO1+*Al*ARC1+*Al*SRK_b_+*Al*SCR_b_ line 81 had 80.2 ± 12.2 pollen grains on average per stigma. Therefore, the Sha wild-type pollen grains were equivalent to those of the Sha *Bn*GLO1+*Al*ARC1+ *Al*SRK_b_+*Al*SCR_b_ lines (group a) when analyzed by a Welch’s ANOVA followed by a Levine test, while the self-incompatible lines of Sha *Al*ARC1+*Al*SRK+*Al*SCR were all similar (group b) as summarized in Figure 3B (P<0.05). These results show that the Sha *Bn*GLO1+*Al*ARC1+ *Al*SRK_b_+*Al*SCR_b_ lines are much closer to Sha wild-type pollen grain adherence implying a loss of the ability to reject self-pollen grains at very early stages of the pollen-pistil interaction.

### Self-incompatible lines expressing *Bn*GLO1 had higher levels of seed set

Since the self-incompatible lines that were expressing *Bn*GLO1 had similar amounts of pollen grains that were able to adhere to the stigma surface 2 hours post pollination, we were curious to ask if the expression of *Bn*GLO1 lead to increased seed set. Therefore, plants were manually self-pollinated and allowed to develop. Mature siliques were dissected 10 days post-pollination and the total number of seeds per silique were tallied (Figure 4). Sha wild-type plants had an average of 50.2 ± 2.3 seeds per silique, while in contrast the Sha *Al*ARC1+*Al*SRK_b_+*Al*SCR_b_ lines showed much lower number of seeds; line 1 had 9.8 ± 6.8 seeds per silique, line 2 had high variance with 20.1 ± 13.1 seeds per silique, while line 5 had 18.1 ± 5.2 seeds per silique. The Sha *Bn*GLO1+*Al*ARC1+*Al*SRK_b_+*Al*SCR_b_ lines had much higher numbers of seeds per silique; line 61 had 42.1 ± 6.3 seeds per silique, line 76 had 40.6 ± 5.4 seeds per silique and line 81 had 28.0 ± 5.3 seeds per silique.

**Figure 4.**
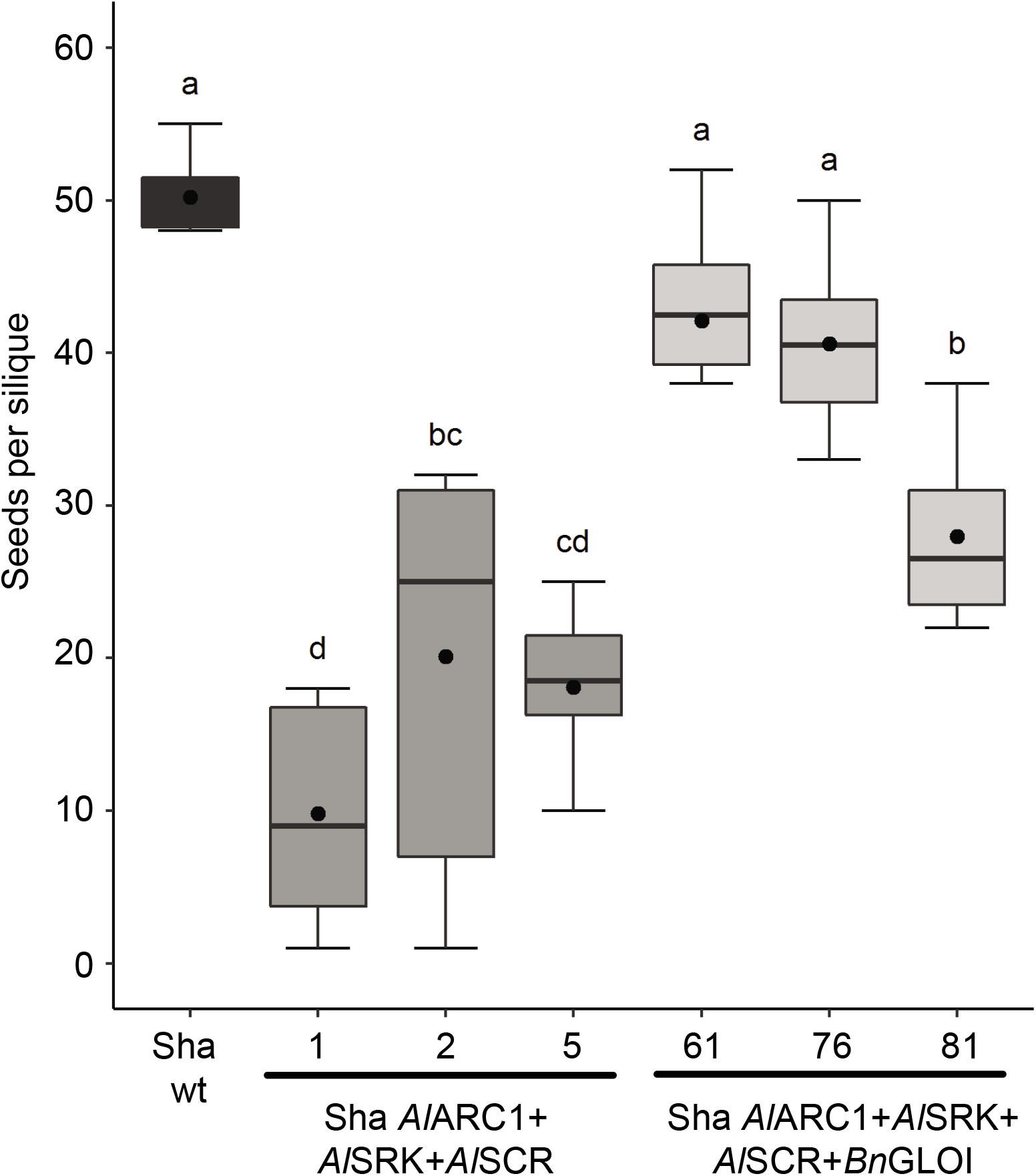
Overexpression of BnGLO1 in self-incompatible pistils restores the acceptance of incompatible pollen grains resulting in seed set. To determine if the expression of *BnGLO1* in self-incompatible pistils had a long-term impact on pollen-pistil interactions, flowers were manually self-pollinated and allowed to set seed. 10 to 11 days after manual pollination, siliques were dissected, and the total number of seeds tallied. These data were analyzed with a one-way ANOVA with a Tukey post hoc test (p < 0.5), n = 10 siliques per each transgenic line. Relative Humidity was between 35-55% during manual pollinations and subsequent time in the growth chambers.

To determine the significance of these results, these data were examined with an ANOVA followed by a Tukey test (P<0.05), letters indicate lines that are statically similar e.g. Sha wild-type, Sha *Bn*GLO1+*Al*ARC1+*Al*SRK_b_+*Al*SCR_b_ line 61 and Sha *Bn*GLO1+*Al*ARC1+*Al*SRK_b_+*Al*SCR_b_ line 76. The self-incompatible lines of Sha *Al*ARC1+*Al*SRK_b_+*Al*SCR_b_ line 1 and Sha *Al*ARC1+ *Al*SRK_b_+*Al*SCR_b_ line 5 were also statistically similar with overlap between line 5 and Sha *Al*ARC1+*Al*SRK_b_+*Al*SCR_b_ line 2. Interestingly, the self-incompatible line Sha *Al*ARC1+ *Al*SRK_b_+*Al*SCR_b_ line 2 was determined to be similar to that of Sha *Bn*GLO1+*Al*ARC1+ *Al*SRK_b_+*Al*SCR_b_ line 81. These data seem to indicate that the ability of some pollen grains to adhere to the stigma surface is strong enough to overcome the self-incompatibility response, even in the Sha *Al*ARC1+*Al*SRK_b_+*Al*SCR_b_ line 2. Overall, two of the three *Bn*GLO1 lines were statistically similar to wildtype, thus demonstrating a breakdown or overcoming of the selfincompatibility response.

## Discussion

Our results demonstrate that the role of GLO1 is conserved between pollen acceptance in *B. napus* and artificial self-incompatible *A. thaliana*. By using well characterized self-incompatible *A. thaliana* (Indriolo et al. 2014), the addition of *Bn*GLO1 was demonstrated to break down the self-incompatible pollen response in *A. thaliana*. The relationship of *Bn*GLO1 to glyoxalases in *A. thaliana* and *A. lyrata* was determined using an alignment and generation of a phylogenetic tree based on the full-length protein sequences of the ten most closely related glyoxalases from both *Arabidopsis* species. The phylogenetic tree revealed that a single copy from both *A. thaliana* and *A. lyrata* formed a clade with *Bn*GLO1 and that when these protein sequences were aligned, they were the most highly conserved. Furthermore, *At1g11840*, the most closely related *A. thaliana* glyoxalase is expressed in the stigma along with other tissues, implying that it may play a role in the stigma during pollinations.

The confocal imaging of unpollinated stigmas demonstrated that *Bn*GLO1-RFP construct localized to the cytoplasm, as was observed when it was expressed in *B. napus* (Sankaranarayanan et al. 2015). We qualitatively assessed the phenotype of the plants using aniline blue staining to examine self-pollinations and the artificial self-incompatible lines expressing *Bn*GLO1 clearly accepted ‘self’ pollen grains. Quantitative analysis of pollen grain adherence demonstrated that the *Bn*GLO1 expressing self-incompatible lines were statistically similar to wildtype via the ANOVA, and that the total number of pollen grains was between 79.1-86.2% of what was observed for wildtype (=100%). Therefore, within the first 2 hours following pollination, the *Bn*GLO1 expressing plants are very similar to wildtype and quite different than the self-incompatible lines which showed between 18.8-30.7% adherence compared to wildtype.

The seed set results overall contribute to the hypothesis that expression of *Bn*GLO1 breaks down the self-incompatible pollination response, with lines 61 and 76 being statistically similar to that of wildtype seed set and had around 83-85% of the total seed set of wildtype (=100%). However, line 81 had a much stronger self-pollen rejection phenotype and was statistically similar to a single self-incompatible line, number 2 having only 56% of the amount of seeds per silique as wildtype. This amount of variance between transgenic lines is not unusual though; when *Bn*GLO1-RFP was overexpressed in a *B. napus* W1 (self-incompatible) background, the seed set was not uniform between the three independent lines. One line had an average seed set of 6.5 while the other two were between 1-1.5 seeds per pod and a compatible line had an average of around 29 seeds per pod (Sankaranarayanan et al. 2015). When examined as the percent of a compatible pollination (where 29 seeds = 100%) the *Bn*GLO1-RFP overexpression lines were between 3-22% of wildtype when compared to the seed set of 0 seeds per pod in a W1 self-incompatible pollination (Sankaranarayanan et al. 2015). Considering the behavior of *Bn*GLO1-RFP in *B. napus* compared to artificial self-incompatible *A. thaliana*, we are not surprised that these are not clear-cut phenotypes; instead these experiments show variation between the phenotypes of different lines. Some of that that variation applies to the fact that the pollen grain adherence of Sha *Bn*GLO1+*Al*ARC1+*Al*SRK_b_+*Al*SCR_b_ line 81 had approximately 81 pollen grains per stigma and was higher than that of Sha *Al*ARC1+*Al*SRK_b_+*Al*SCR_b_ line 2 that had 31.3 pollen grains per stigma, but that pollen grain adherence between these lines did not correspond to the seed set that was observed. Therefore, there may be some additional differences between the artificial self-incompatible lines and our *Bn*GLO1 expressing lines between pollen grain adherence and successful seed set.

Our results indicate that for *Bn*GLO1 to break down the artificial self-incompatible response in *A. thaliana*, there is a dose dependent aspect of *Bn*GLO1 in this process. Recent studies have highlighted how the whole genome triplication in *Brassica* has impacted the function of the regulatory genes in the self-incompatibility pathway in *Brassica* species. After the whole genome triplication event, most of the self-incompatible signaling pathway genes were maintained as single copy genes in diploid *Brassica* species. These genes included; *ARC1, JDP1, THL1, THL2, Exo70A1*, while *PLD1* and *GLO1* were kept as duplicated genes (Azibi et al. 2020). Further analysis of *GLO1* showed that there may be an expression dosage difference involved in part between self-incompatible and self-compatible *B. rapa* cultivars. The self-compatible *B. rapa* var. *trilocularis*, ‘Z1’ line when examined for *BnGLO1* expression showed expression of the *GLO1-MF1* (A08) and *GLO1-MF2* (A06) genes in stigma cDNA samples. These results show that there are two copies of *GLO1* expressed in the stigma of self-compatible *B. rapa*. In contrast, the *B. rapa* var. *pekinensis*, ‘Chiifu’ is a self-incompatible line and only the *GLO1-MF1* (A08) copy of *GLO1* is expressed. Granted the relative expression level is not equal in the self-compatible ‘Z1’, between the *GLO1-MF1* and *GLO1-MF2* in that background but it is present versus absent in the ‘Chiifu’ *B. rapa* (Azibi et al. 2020). Our use of expressing an extra copy of *BnGLO1* in our self-incompatible *A. thaliana* background may be similar to what is happening between these two different *B. rapa* varieties; having more GLO1 activity present in the stigma could help in accepting pollen grains regardless of the activation of the SI response.

These results lead us to propose a model for what is happening in the *Al*ARC1+ *Al*SRK_b_+*Al*SCR_b_ lines compared to the *Bn*GLO1+*Al*ARC1+*Al*SRK_b_+*Al*SCR_b_. In our artificial self-incompatible Sha *A. thaliana* (Figure 5A), the pollen ligand *Al*SCR_b_ is recognized by *Al*SRK_b_. As *Al*SRK_b_ is phosphorylated and activated, it in turn activates *Al*ARC1 which targets downstream factors including *At*GLX1 for degradation by ubiquitination (Sankaranarayanan et al. 2017; Schmitz et al. 2018). The *At*GLX1 levels then decrease and more proteins become MG-modified helping in the self-incompatible pollen response. In contrast, in our *Bn*GLO1+*Al*ARC1+*Al*SRK_b_+*Al*SCR_b_ plants (Figure 5B), the self-incompatibility pathway is still activated, but the level of glyoxalase I is much higher than in a normal stigma with the additional expression of the *Bn*GLO1-RFP. Despite *Al*ARC1 being able to ubiquitinate protein targets, there are higher levels of *At*GLX1 and *Bn*GLO1-RFP present and they are able to overcome the activity of *Al*ARC1, detoxifying MG-modification of proteins and allowing the stigma to accept the ‘self’ pollen.

**Figure 5.**
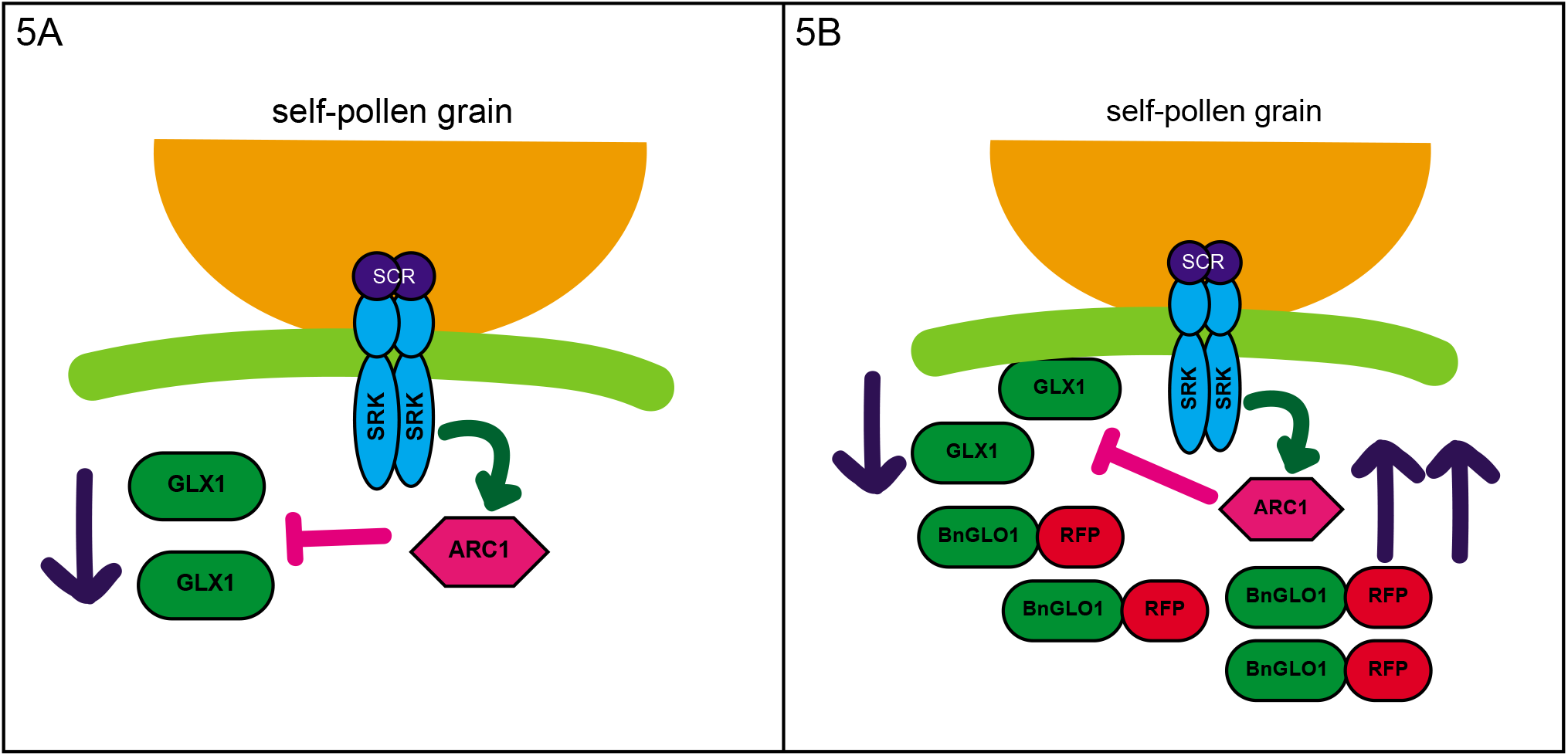
Model of BnGLO1 action during self-incompatible pollinations. 5A.) During a self-pollination in the artificial self-incompatible *A. thaliana*, SRK is able to detect SCR on the pollen grain surface. SRK becomes phosphorylated and activated, and in turn phosphorylates ARC1. ARC1 then ubiquitinates GLX1 which decreases due to degradation in the 26S proteasome. 5B.) With the addition of *Bn*GLO1-RFP in the artificial self-incompatible *A. thaliana*, there is more glyoxalase present than normal. Even though SCR is still able to activate SRK and ARC1 activity, the level of GLX1 and *Bn*GLO1-RFP is high enough that ARC1 cannot ubiquitinate all of it for degradation in the 26S proteasome. The higher amounts of GLX1 and *Bn*GLO1 then facilitates the acceptance of the ‘self’ pollen grain.

Artificial self-incompatibility in *A. thaliana* is an ideal system to study and validate compatibility and self-incompatibility genes identified in other related species. *Bn*SCR1 or *Bn*SRK1 have been previously shown to be non-functional in *A. thaliana*, as expression of *Bn*SCR1-*Bn*SRK1-*Bn*ARC1 failed to reconstitute self-incompatibility in self-pollinated *A. thaliana* (Zhang et al. 2019). There are several pieces of evidence that support the idea of differences between *Brassica* and *Arabidopsis* self-incompatibility signaling pathways. These observed differences arise due to variations in the upstream components of self-incompatibility signaling (S-haplotype genes) and the downstream components of the signaling pathway like ARC1 and interacting partners including EXO70A1, GLO1 and PLD1 are conserved. Previously it has been demonstrated that expression of RFP: *Bn*Exo70A1 is able to rescue the stigmatic fertility defect observed in *A. thaliana exo70A1* mutants suggesting that there is an evolutionarily conserved function for Exo70A1 in the dry stigmas of the *Brassicaceae* (Samuel et al. 2009).

Despite the divergence of *Brassica* from *Arabidopsis* some 20 million years ago, *Bn*GLO1 shares a high sequence similarity to the stigma expressed glyoxalase I homologs in *Arabidopsis thaliana* (Franzke et al. 2011; Wang et al. 2011). It is not surprising that *Bn*GLO1 could function in *A*. *thaliana* unlike BnSCR1 and *Bn*SRK1 that show a high degree of variation in their sequences. It would be interesting to see if a reciprocal experiment of transferring *Arabidopsis GLX1* genes into self-incompatible *B*. *napus* lines could rescue the SI phenotype. Presence of a large GLX1 gene family (11 members) in *A. thaliana* and a high functional redundancy makes it difficult to identify their plausible role in pollination as single mutants often fail to exhibit any pollination or fertility phenotypes. Generation of higher order knockout mutants of GLX1 homologues in *A. thaliana* or stigma specific suppression of multiple *At*GLX1s using an RNAi approach is required to unequivocally demonstrate the role of GLX1 in mediating compatible pollination in *Arabidopsis*. Nevertheless, in this study we were able to circumvent the problem of functional redundancy and unmask the functional role of GLO1 in *A. thaliana* pollination using the artificial self-incompatibility system. Phospholipase D (PLD) is another compatibility factor that exists as a part of a large gene family in *Arabidopsis*; the same approach can be used to validate its role in pollination by utilizing the *A. thaliana* artificial self-incompatibility system. These proposed experiments will further help us to understand the conservation of self-incompatibility signaling mechanisms and components in *Arabidopsis* and *Brassica*. In conclusion, this study has validated the functional role of GLO1 as a compatibility factor in *Arabidopsis* self-incompatibility and our study supports the idea that despite the divergence of upstream self-incompatibility signaling components of *Arabidopsis* and *Brassica*, the downstream signaling components are conserved between the two species.

## Contributions

SS and EI initiated the project and designed research; PK, SS, MB and EI performed all of the experiments. MB and EI statistically analyzed the results. EI and SS wrote the manuscript. EI obtained funding and supervised the project. All authors discussed the results and commented on the manuscript. The authors would lastly like to thank Ms. Alejandra Cobos and Dr. Marcus Samuel for critical feedback on the manuscript.

---

***Supplemental Figure 1. Additional Aniline Blue stains of A. thaliana stigmas 2 hours post pollination*.**

Manual self-pollination results of Sha wild-type compatible pollinations, Sha incompatible pollinations of *AlARC1 +AlSRK_b_+AlSCR_b_* lines and the Sha *BnGLO1+AlARC1+AlSRK_b_+AlSCR_b_* lines. Scale bar = 50 μm.

***Supplemental Figure 2*.** The reproductive expression data series of *Arabidopsis thaliana Glyoxalase* family members generated with Arabidopsis Heat Tree Viewer.

***Supplemental Figure 3. PCR genotyping of all transgenic A. thaliana lines*.**

*A. thaliana* Sha plants were genotyped for the presence of transgenes. Line 61, 76 and 81 were the plants that were positive for the presence of *AlARC1, AlSCR-AlSRK* and *BnGLO1* with primers that were specific for each of these constructs. Sizes for each PCR product are included in the figure. The previously characterized *A. thaliana* Sha artificial self-incompatible lines 1, 2 and 5 were confirmed by PCR with the *AlARC1* and the *AlSCR-SRK* constructs.

***Supplemental Figure 4. R script*.**

R script used to analyze seed set and pollen grain adherence.

***Table 1. Primers used in genotyping transgenic plants***.

