## Supplementary figures and images for "Expression of *Brassica napus* GLO1 is sufficient to breakdown artificial self-incompatibility in *Arabidopsis thaliana*"

### Supplemental Figure 1

Supplemental Figure 1

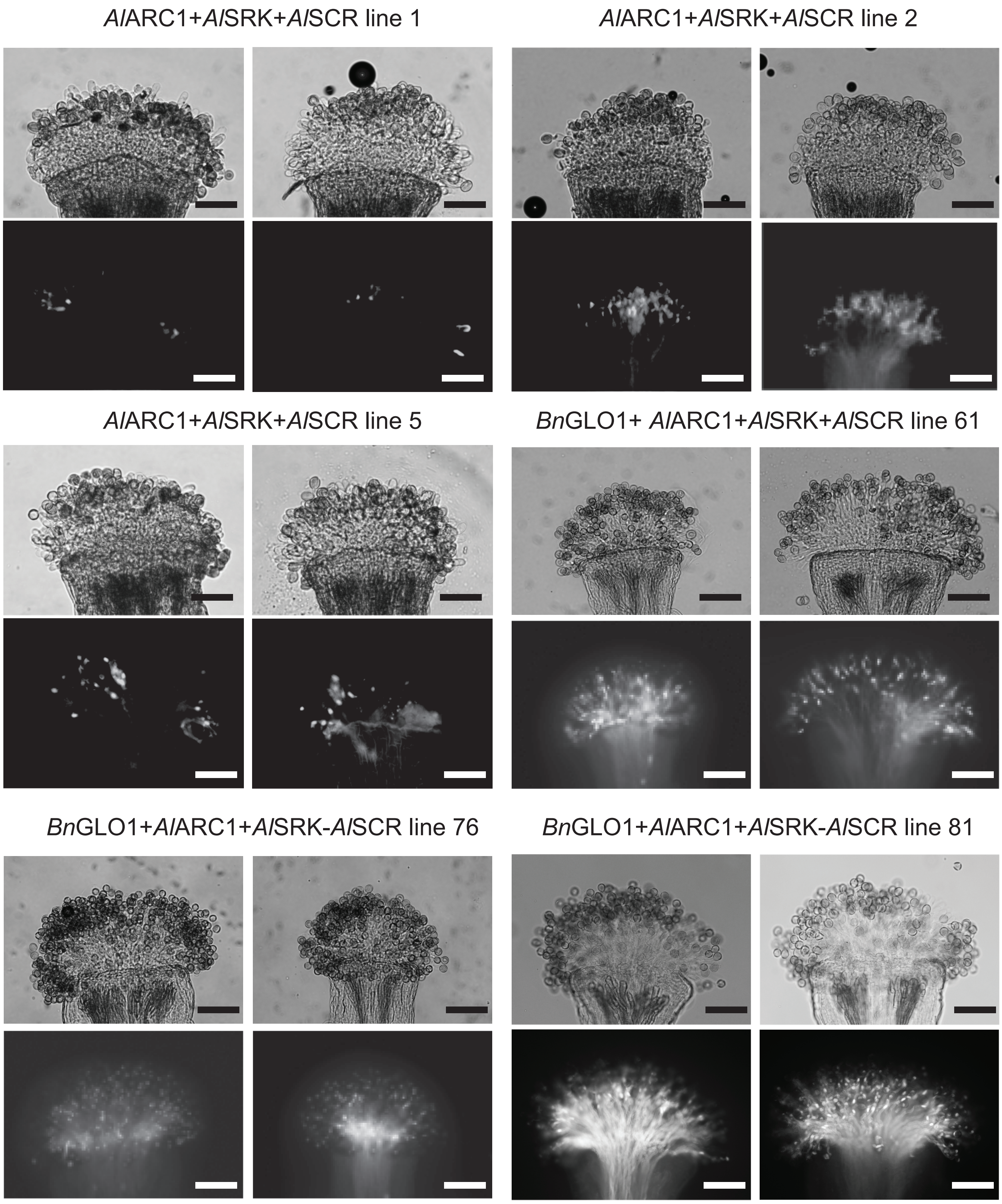
