## Supplemental Figure 2 for "Expression of *Brassica napus* GLO1 is sufficient to breakdown artificial self-incompatibility in *Arabidopsis thaliana*"

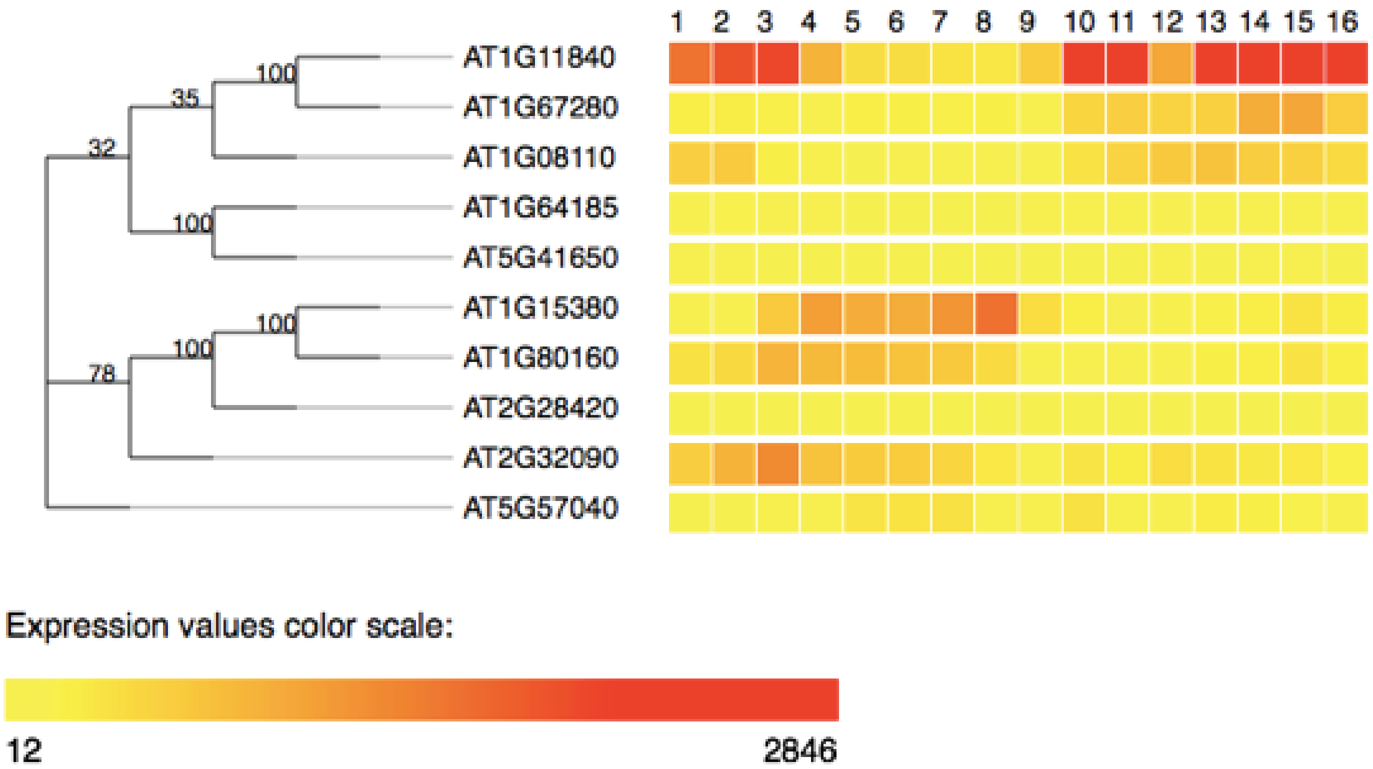


Selected expression profiles:

| | |
|---|---|
| 1 | Uninucleate Microspore (H) |
| 2 | Bicellular Pollen (H) |
| 3 | Tricellular Pollen (H) |
| 4 | Mature Pollen Grain (H) |
| 5 | Dry Pollen (Q) |
| 6 | .5h Pollen Tubes (Q) |
| 7 | 4h Pollen Tubes (Q) |
| 8 | Semi-in-Vivo Pollen Tubes (Q) |
| 9 | Sperm (Bor) |
| 10 | Stigma (S) |
| 11 | Ovary (S) |
| 12 | Ovules (Boa) |
| 13 | Unpollinated Pistil (Boa) |
| 14 | Pistil 0.5h after Pollination (Boa) |
| 15 | Pistil 3.5h after Pollination (Boa) |
| 16 | Pistil 8h after Pollination (Boa) |
