## Supplemental Figure 3 for "Expression of *Brassica napus* GLO1 is sufficient to breakdown artificial self-incompatibility in *Arabidopsis thaliana*"

*A. thaliana* Sha plants with *Bn*GLO1, *Al*SCR-*Al*SRK & *Al*ARC1 transgenes

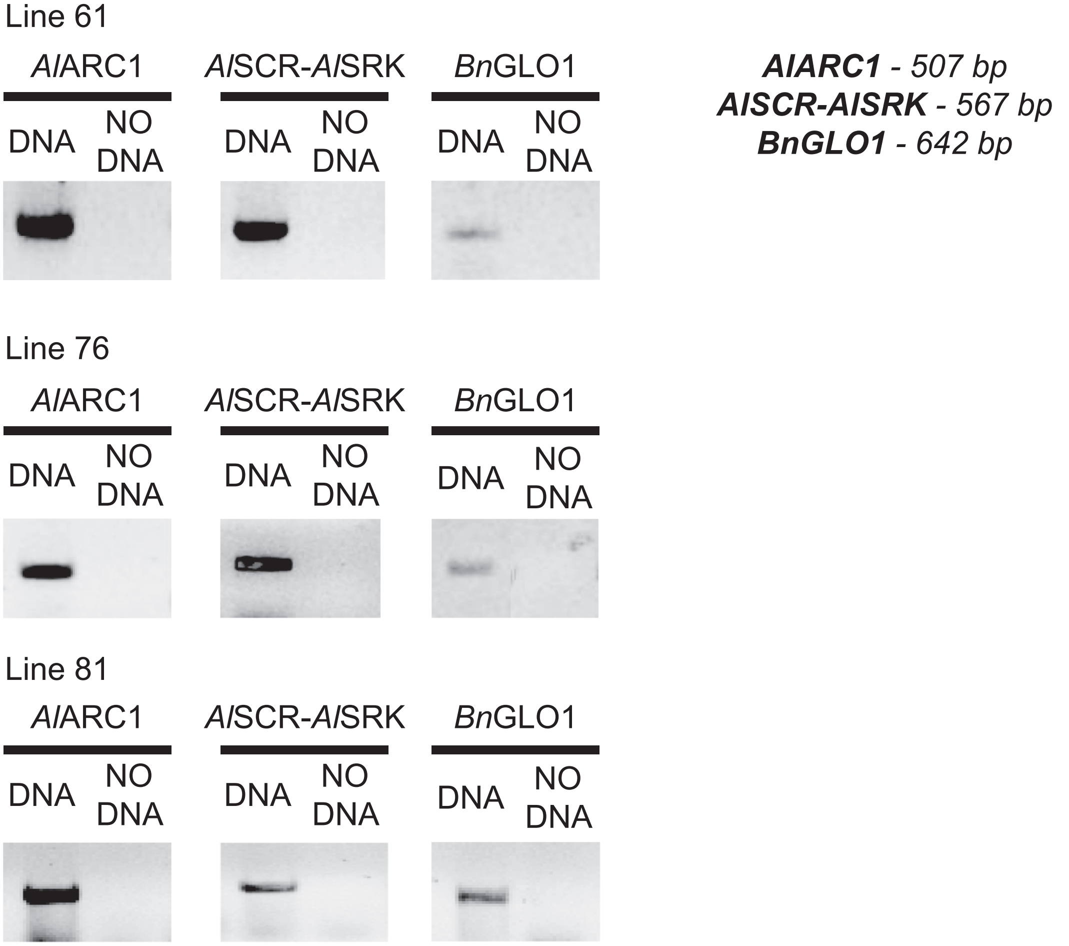


*A. thaliana* Sha plants with *Al*SCR-*Al*SRK & *Al*ARC1 transgenes

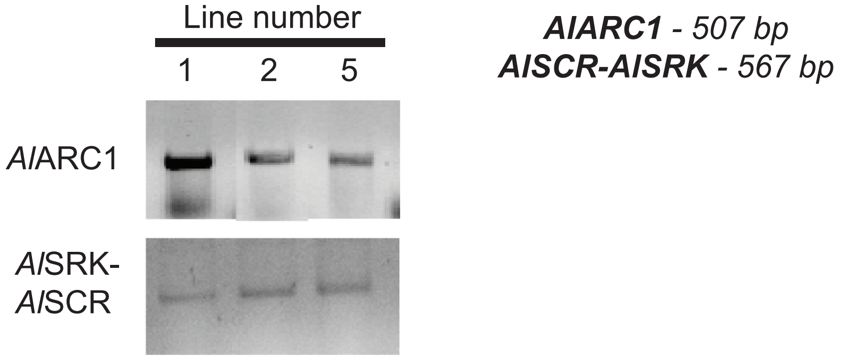
