## Supplemental Figure 4 for "Expression of *Brassica napus* GLO1 is sufficient to breakdown artificial self-incompatibility in *Arabidopsis thaliana*"

```
seedset <- read.csv("seedset.csv")
library(ggplot2)
install.packages("sf", dependencies = TRUE)
install.packages("Matrix", dependencies = TRUE)
install.packages("agricolae", dependencies = TRUE)
library(agricolae)
library(tidyverse)
install.packages("dplyr", dependencies = TRUE)
library(dplyr)


abs_max <- max(seedset$seed.set)
maxs <- seedset %>%
  group_by(line) %>%
  summarise(seed.set=max(seed.set) + 0.05 * abs_max)


Tukey_test <- aov(seed.set ~ line, data=seedset) %>%
  HSD.test("line", group=TRUE) %>%
  .$groups %>%
  as_tibble(rownames="line") %>%
  rename("Letters_Tukey"="groups") %>%
  select(-seed.set) %>%
  left_join(maxs, by="line")
```

```
tiff("Sha BnGLO1 seedset.tiff", units="in", width=4.5, height=4.5, res=300)
ggplot(seedset, aes(line, seed.set)) +
  stat_boxplot( aes(line, seed.set),
                geom='errorbar', linetype=1, width=0.5)+
  geom_boxplot( aes(line, seed.set),outlier.shape=NA) +
  stat_summary(fun.y=mean, geom="point", size=5) +
  geom_text(data=Tukey_test, aes(label=Letters_Tukey), size = 4) +
  labs(y="Number of Seeds per Silique", face="bold", size="16") +
  scale_fill_grey() +
  theme_classic() +
  theme(axis.title.x=element_blank()) +
  theme(axis.text.x = element_blank()) +
  theme(axis.title = element_text( size = 9)) +
  theme(legend.position = "none") +
  coord_cartesian(ylim=c(0, 62)) +
  scale_y_continuous(breaks=seq(0, 60, 10))

dev.off()
```
