## Supplementary material for "Expression of *Brassica napus* GLO1 is sufficient to breakdown artificial self-incompatibility in *Arabidopsis thaliana*": Table 1

| Gene | direction | sequence | purpose |
| --- | --- | --- | --- |
| AlSCRb | left primer | AACAAGTGCATGCGTTCTGA | genotyping |
| AlSCRb | right primer | ACAGTCGCATAAACGTGCAA | genotyping |
| AlSRKb | left primer | GTGGTGGCAGAGCTTCTTTC | genotyping |
| AlSRKb | right primer | ACAAATCGGTGACGTGTTCA | genotyping |
| AlARC1 | left primer | CAAATGTAGATGGCTGCATCA | genotyping |
| AlARC1 | right primer | ACATCATCGCTGTGTTTTCG | genotyping |
| BnGLO1 | left primer | GCTGATTTGGTGGAGTGGCC | genotyping |
| BnGLO1 | right primer | CCTCATTCCAGTTCCTTCAG | genotyping |
